# LATEER: Low-Cost Open-Source Platform for Electrical Stimulation and TEER Measurement in Human Cardiomyocytes

**DOI:** 10.64898/2026.08.06.743263

**Authors:** Florencia De Lillo, Joaquin Smucler

## Abstract

Electrical stimulation (ES) and transepithelial/transendothelial electrical resistance (TEER) measurements are essential techniques in cell biology and tissue engineering, yet commercial devices for these applications cost between USD 2,500–9,000 and typically offer only one functionality. We present LATEER (Low-cost Arduino-based TEER and Electrical stimulation device), an open-source hardware platform that combines both ES and TEER measurement capabilities at a total cost below USD 100. The device features four independent channels, configurable pulsatile signals (amplitude up to 8.2 V, frequency 0.1–500 Hz, pulse width ≥0.1 ms), and a resistance measurement range of 300 Ω to 1 MΩ, with <5% error for R □ 4.7 kΩ. LATEER uses commercially available graphite pencil leads as electrodes (∼USD 2 vs. USD 350 for commercial Ag/AgCl electrodes), which demonstrated excellent biocompatibility in cell culture. The system includes 3D-printed electrode holders compatible with standard 12-well and 24-well plates, allowing microscope visualization without electrode removal, and a Python-based graphical user interface for parameter configuration and real-time data acquisition. Because the electrodes remain fixed in the plate lid and only a single cable enters the incubator, both stimulation and resistance measurement can run continuously under standard culture conditions (37 °C, 5% CO□) without removing the plate or repositioning the electrodes, avoiding the temperature excursions and placement variability inherent to manual chopstick measurements. Validation with human pluripotent stem cell-derived cardiomyocytes demonstrated reliable frequency capture (electrical pacing) of the contracting monolayer, with a capture threshold between 250 and 400 mV/mm and controlled pacing across the 0.5–5 Hz range. TEER functionality was verified with mesenchymal stem cells, where the device resolved cell-density-dependent differences in electrical resistance in real time. All design files, firmware, and software are freely available under the CERN-OHL-S v2 license, enabling replication and customization by research laboratories worldwide.

**Highlights:**

- An open-source device combines electrical stimulation and TEER measurement under $100
- Graphite electrodes offer biocompatibility at 0.6% cost of commercial alternatives
- Four independent channels with configurable parameters and real-time data logging.
- Continuous run setup in-incubator; no electrode repositioning needed
- Validated with stem cell-derived cardiomyocytes, achieving frequency capture (threshold 250–400 mV/mm)
- 3D-printed holders enable microscope visualization without electrode removal

**Graphical abstract:** 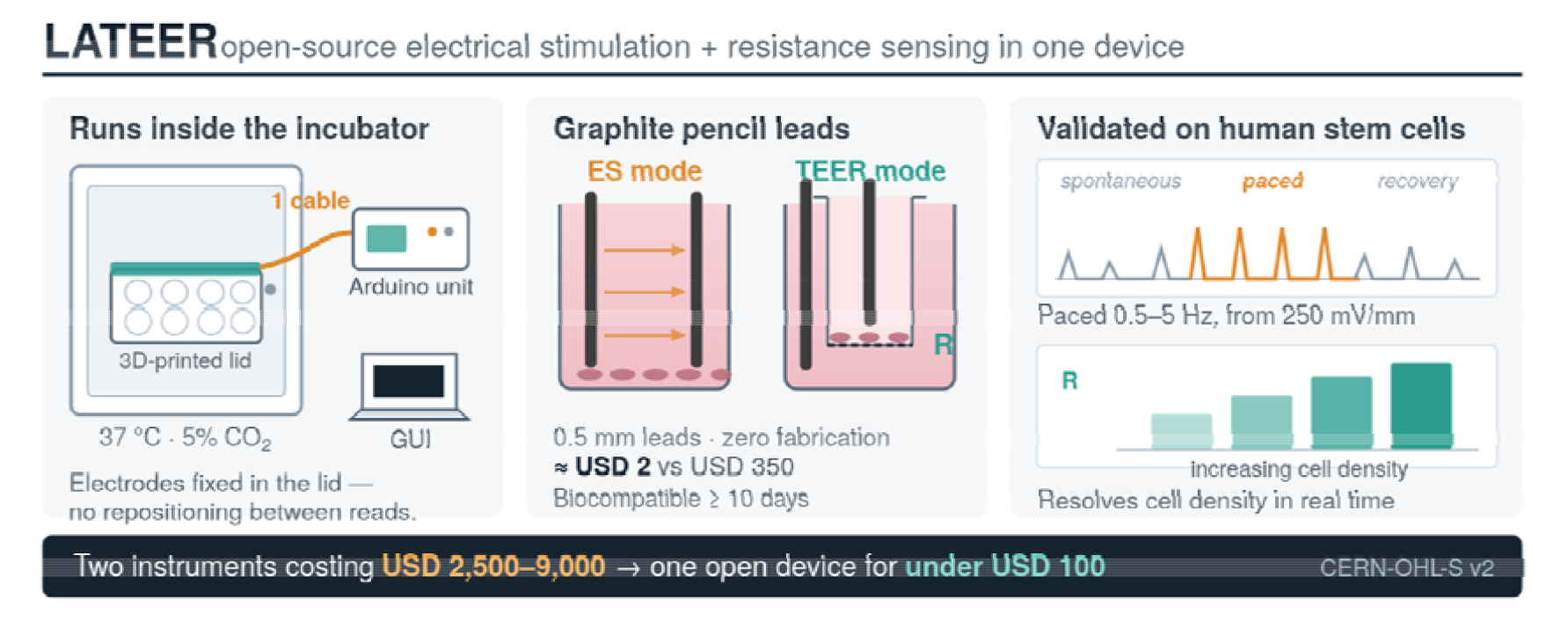

**Specifications Table:** 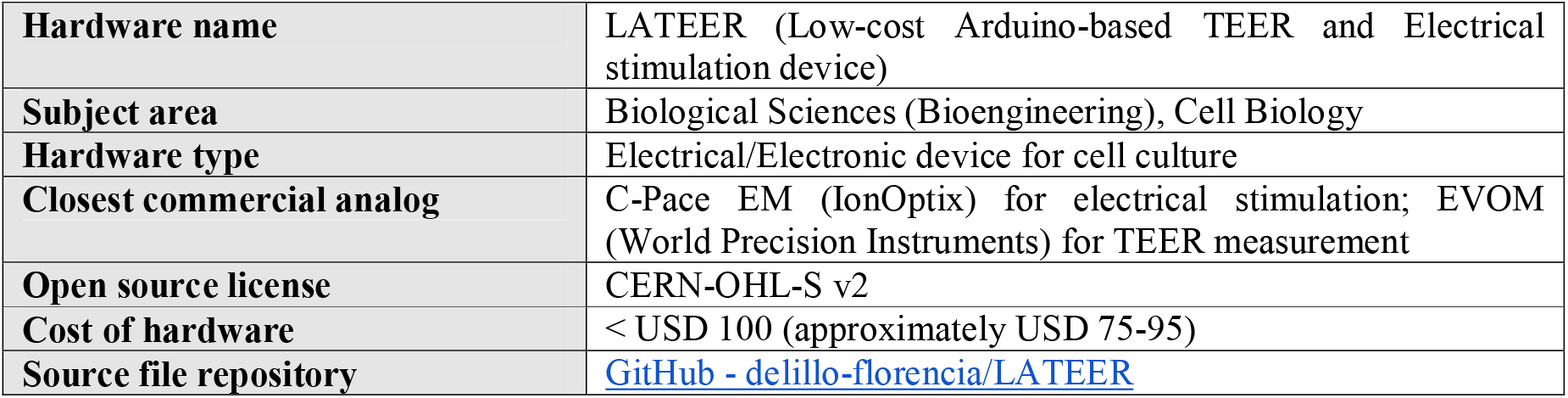

## Hardware in context

Electrical stimulation (ES) is a versatile technique in tissue engineering that influences fundamental cellular processes, including proliferation, migration, alignment, and differentiation across multiple cell types ^1^. ES has been applied to promote osteogenic differentiation of mesenchymal stem cells ^2^, enhance neural stem cell differentiation and neurite outgrowth ^3^, and induce alignment and gene expression changes in various excitable and non-excitable cells ^4^. In cardiac tissue engineering, ES has become a critical tool for promoting maturation of human embryonic stem cell-derived cardiomyocytes (hESC-CMs), which typically exhibit immature, fetal-like phenotypes including disorganized sarcomeres, absence of T-tubules, and spontaneous rather than stimulus-triggered contractions ^5^. Application of pulsed electrical fields has been shown to improve sarcomere organization, enhance calcium handling, increase expression of cardiac-specific genes and ion channels, and promote cellular elongation and alignment ^6–8^.

Transepithelial/transendothelial electrical resistance (TEER) is a non-invasive technique that measures the electrical impedance across cellular monolayers, reflecting the ionic conductance of the paracellular pathway ^9^. While predominantly used to assess tight junction integrity and barrier function in epithelial and endothelial models—including blood-brain barrier, gastrointestinal tract, and pulmonary systems—impedance-based approaches can also provide information about membrane capacitance, which correlates with cell membrane area and differentiation state ^10^.

Commercial devices for ES, such as the C-Pace EM (IonOptix), cost approximately USD 9,000, while TEER measurement systems like the EVOM (World Precision Instruments) cost around USD 2,500, with additional electrode costs of USD 350 per unit. These high costs represent a significant barrier for many research laboratories, particularly in developing countries.

Table 1 and Table 2 summarize how LATEER compares with the two most widely used commercial references, the EVOM (TEER) and the C-Pace EM (ES). LATEER reproduces the core functionality of both at a fraction of the cost, and adds continuous in-incubator operation and a multi-user graphical interface. Its main present limitation relative to the commercial units is the use of direct rather than alternating current, addressed in the future-work section.

**Table 1.** Comparison of LATEER with the EVOM TEER meter.

| Parameter | EVOM | LATEER |
| --- | --- | --- |
| Signal | Alternating current | Direct current (pulsed) |
| User electrode setup required | Yes | No |
| Electrode material | Ag/AgCl | Graphite |
| Continuous in-incubator measurement | No | Yes |
| Electrode cost | USD 350 | USD 2 |
| System cost | USD 2,500 | < USD 100 |

**Table 2.** Comparison of LATEER with the C-Pace EM electrical stimulator.

| Parameter | C-Pace EM | LATEER |
| --- | --- | --- |
| Amplitude | $\pm 40$ V | +8.2 V |
| Pulse width | 0.4–24 ms | 0.1–500 ms |
| Frequency | 0.01–99 Hz | 0.1–500 Hz |
| Graphical user interface | Basic | Yes |
| Electrical stimulation | Yes | Yes |
| System cost | USD 9,000 | < USD 100 |

Several research groups have developed custom solutions to address these limitations. Jones and Chen ^11^ reported an Arduino-based TEER measurement system using stainless steel electrodes coated with carbon ink. Raut et al. ^12^ developed an automated TEER system with 3D-printed electrode holders, though still using commercial STX2 electrodes (∼USD 350 each). More recent open-source TEER instruments have refined this approach; Lattanzi et al. reported an Arduino-based meter that, like the present work, avoids a continuous same-polarity bias to limit electrode double-layer formation and cell damage ^13^. In parallel, several open-source electrical-stimulation platforms for hESC-derived cardiomyocytes have appeared, including the electromechanical BEaTS-α device ^14^, the low-cost cardiac-stimulation bioreactor of Licata et al. ^15^, and a dynamic (time-varying frequency/pulse-duration) stimulation device for hESC-CM differentiation reported by Kalkunte et al. ^16^, which, notably, also uses graphite (carbon-block) electrodes, cast in PDMS rather than off-the-shelf pencil leads. Dual-function instruments that combine stimulation with impedance/TEER sensing have also been demonstrated, notably the ECSARA apparatus ^17^. Taken together, the landscape as of 2026 offers capable open-source instruments for each function separately — TEER meters ^11,13,18^ and stimulators ^14,15,19,20^ — but the one dual-modality instrument is custom, closed and comparatively costly. Although Arduino-based resistance measurement, ES+TEER integration and the use of carbon electrodes for cell stimulation have all been reported previously, no single, fully open, sub-USD-100 unit currently can perform both functions on standard multiwell plates using electrodes that require no fabrication step. LATEER is presented as that combination.

Here we present LATEER (Low-cost Arduino-based TEER and Electrical stimulation device), an open-source hardware platform that provides both ES and TEER measurement in a single unit — each function selected per experiment by fitting the corresponding electrode lid **—** at a total cost below USD 100 . The device uses commercially available graphite pencil leads as electrodes, which offer excellent biocompatibility at minimal cost (∼USD 2 vs. USD 350 for commercial electrodes). The system includes:

- Four independent channels for electrical stimulation or TEER measurement
- Configurable pulsatile signals with amplitude up to 8.2 V, frequency 0.1-500 Hz, and pulse width ≥0.1 ms
- Resistance measurement range of 300 Ω to 1 MΩ (<5% error for ⊡ l-4.7 kΩ)
- 3D-printed electrode holders compatible with 12-well and 24-well culture plates
- Python-based graphical user interface (GUI) for parameter configuration and data acquisition
- Design that allows microscope visualization without removing electrodes

## Hardware description

LATEER consists of four main components: (1) graphite electrodes, (2) 3D-printed electrode holders, (3) an electronic control circuit, and (4) a software interface.

### 2.1 Electrodes

Commercial 0.5 mm diameter, 2H graphite pencil leads were selected as electrodes after evaluating multiple materials including stainless steel with carbon ink coating and carbon rods. Carbon rods were impractical due to their thickness (6 mm) and material shedding. The final selection was based on biocompatibility testing results (see Section 7.1), cost, availability, and dimensional reproducibility.

Graphite pencil leads — widely termed pencil graphite electrodes (PGEs) — are well characterized as inexpensive, disposable, biocompatible electrodes in electrochemistry and biosensing, where lead hardness and surface pre-treatment govern performance ^21^. Different hardness grades (HB, H, 2H, B) and diameters (0.5, 0.7, 1.0 mm) were compared here; differences in monolayer condition were observed between grades, and the 0.5 mm / 2H lead was selected because it released the least particulate material and requires no additional processing.

As previously mentioned, carbon and graphite electrodes for cell stimulation are established prior art, and a concurrently reported device uses graphite in carbon-block form, custom-cut and cast in PDMS, for the same purpose ^16^. The contribution here is therefore not the material but the form factor. To our knowledge, this is the first application of off-the-shelf, unmodified graphite pencil leads as stimulation/pacing electrodes for a cultured cell monolayer. Its dimension is standardised by the manufacturer, requires no cutting, casting, coating or curing step, and can be replaced in seconds by the end user, which is what makes the electrode a genuinely zero-fabrication component of an open hardware build.

Because the resistance channel is a two-electrode divider, the series resistance of the leads themselves is included in every reading; it is common to all channels and to the blank insert and therefore cancels in between-channel comparisons, but it is one reason the absolute readings reported in Section 7.2 are not interchangeable with normalised TEER values.

For TEER measurements, the selection of graphite pencil leads does not impose constraints on electrode geometry. The electrodes can be positioned in the conventional configuration, with one electrode placed above and one below the transwell membrane, enabling standard resistance measurements across the cell monolayer (**Figure 1A)**. This configuration is widely reported in the literature and is directly compatible with commercially available graphite leads.

**Figure 1.**
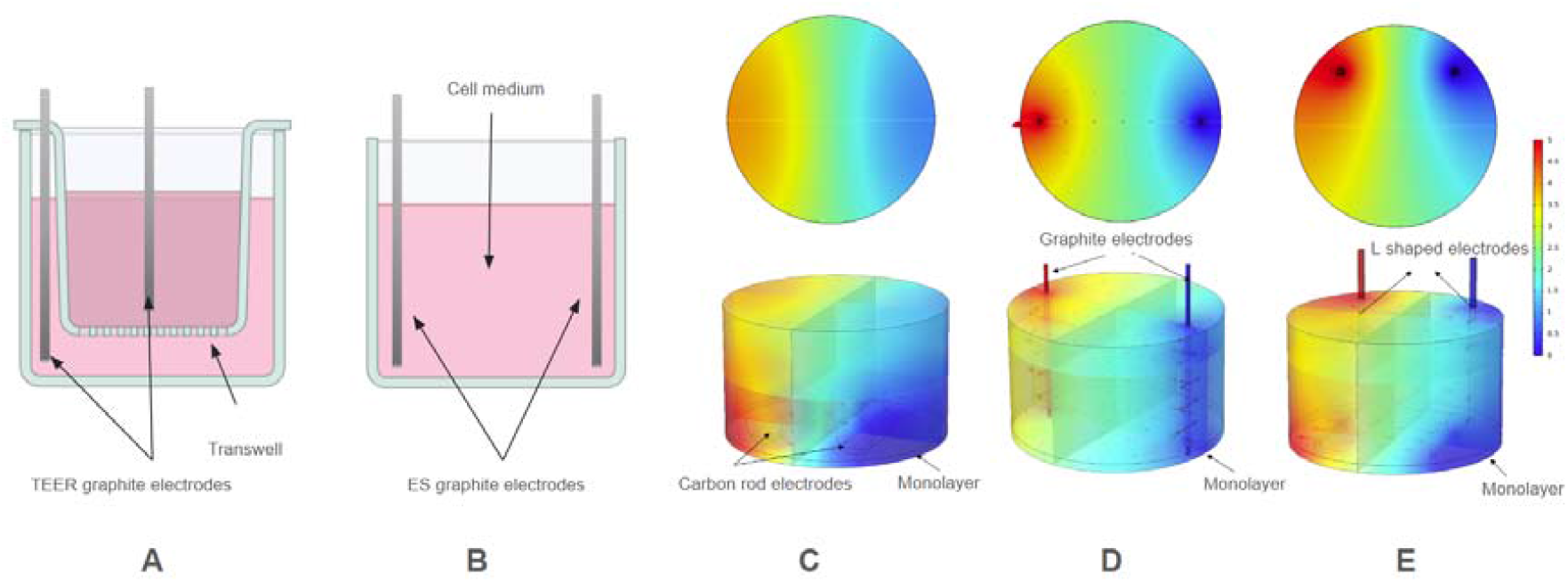
Electrode configurations and electric field simulations. **(A)** Conventional TEER electrode configuration with electrodes positioned above and below the transwell membrane. **(B)** Proposed ES configuration using two vertical, opposed graphite pencil lead as electrodes. **(C–E)** COMSOL Multiphysics® simulations comparing the electric field distributions generated by the vertical graphite electrode configuration, reference L-shaped platinum and carbon rod electrode geometries. While all configurations produce broadly uniform electric fields across the culture area, higher field intensities at the well base are observed for the L-shaped electrodes due to the presence of a horizontal electrode segment near the culture surface.

For ES, however, most reported *in vitro* ES systems employ L-shaped or planar electrodes, with one electrode segment positioned horizontally near the culture surface. These geometries are not compatible with commercially available graphite pencil leads, which are straight and cannot be bent or shaped without fracture. As a result, the use of graphite leads needs an alternative electrode configuration.

To enable electrical stimulation using low-cost graphite electrodes while maintaining compatibility with standard multiwell plates, we implemented a configuration consisting of two straight, vertical electrodes positioned in parallel and opposite to each other within the well **(Figure 1B**). To our knowledge, this electrode configuration using graphite pencil leads has not been previously reported for *in vitro* electrical stimulation of cell monolayers.

COMSOL Multiphysics® simulations were therefore performed to evaluate the electric field distribution produced by the proposed graphite electrode configuration and to compare it against a reference L-shaped platinum and carbon rod electrodes geometry reported in the literature (**Figure 1C-E**). A summary of the simulation setup, including geometry definitions, material properties, and electrostatic modeling assumptions, is provided in **Supplementary Table S1**.

Although all electrode geometries produced a nominally uniform electric field within the bulk culture volume, inspection of the electric field magnitude at the bottom of the well—corresponding to the cell monolayer—revealed marked differences between configurations. In the L-shaped electrode designs, the presence of a horizontal electrode segment positioned close to the culture surface resulted in a locally enhanced electric field at the monolayer, as evidenced by higher |E| values near the well base. In contrast, the vertical graphite electrode configuration produced a more axially distributed field, leading to lower field intensities at the monolayer for the same applied voltage. Consequently, higher applied voltages are required in the graphite configuration to achieve equivalent field strengths at the cellular level. Despite this limitation, the graphite-based configuration was selected due to its substantial advantages in cost, availability, ease of fabrication, and reproducibility, while still operating within physiologically relevant stimulation ranges (validated in Section 7.3).

### 2.2 3D-printed electrode holders

Custom electrode holders were designed in SolidWorks 2019 and manufactured using PLA filament on an Ender 3 Pro 3D printer. A key design feature is the open-ring geometry that allows microscope visualization of cells without removing the electrode assembly, reducing contamination risk during long-term experiments. The holders use a dual metal terminal system for secure graphite lead attachment and easy replacement if breakage occurs **(Figure S1A**,**B)**. Designs are provided for 12-well (MW12) and 24-well (MW24) plates, with parametric files allowing adaptation to other formats. Two holder designs accommodate different experimental configurations: (1) an ES configuration with parallel electrodes at equal height for field stimulation, and (2) a TEER configuration with electrodes positioned above and below the transwell membrane **(Figure S1C**,**D)**.

### 2.3 Electronic circuit

The system is controlled by an Arduino UNO (ATmega328P) microcontroller operating at 5V. The circuit design is based on principles from Adams et al. and Jones and Chen ^11,20^, with modifications to minimize unnecessary cell exposure to voltage.

**TEER measurement circuit:**

Uses a voltage divider principle with a known 320 kΩ reference resistor. Unlike the original design that continuously applies 5V, our implementation uses a PWM-controlled output that only applies voltage during measurement, reducing cell stress. The resistance is calculated using:

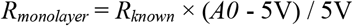

Where A0 is the analog reading. Each measurement averages 10 readings for improved accuracy ^11^.

#### Electrical stimulation circuit

Generates monophasic pulsatile signals using an operational amplifier in non-inverting configuration (gain = 2), a voltage regulator (LM317), and transistor switching. The PWM signal is generated by a digital output of an Arduino microcontroller and fed into the conditioning circuit, where the duty cycle modulates the output amplitude. The pulse frequency and pulse width are controlled by software timing on the Arduino. The circuit provides four independent output channels with individually adjustable amplitudes, while sharing common frequency and pulse width parameters.

### 2.4 Software interface

A graphical user interface (GUI) was developed in Python using PyQt5 and Qt Designer. The interface communicates with the Arduino via USB serial connection and provides: mode selection (ES or TEER), channel configuration (1-4 channels), parameter input for ES (amplitude, frequency, pulse width, duration) or TEER (sampling interval, measurement duration), real-time data visualization, data export to CSV format, and SQL database storage for experiment parameters and results. The interface was designed to be operable by users without a programming or electronics background: all parameters are entered through labeled numeric fields and dropdown menus, wells are selected on an on-screen plate layout mirroring the physical 12-or 24-well plate, and live plots of resistance (TEER mode) or applied waveform (ES mode) update during acquisition **(Figure S2)**. This lowers the practical barrier to routine use of the device in a standard cell-culture lab.

## Design files summary

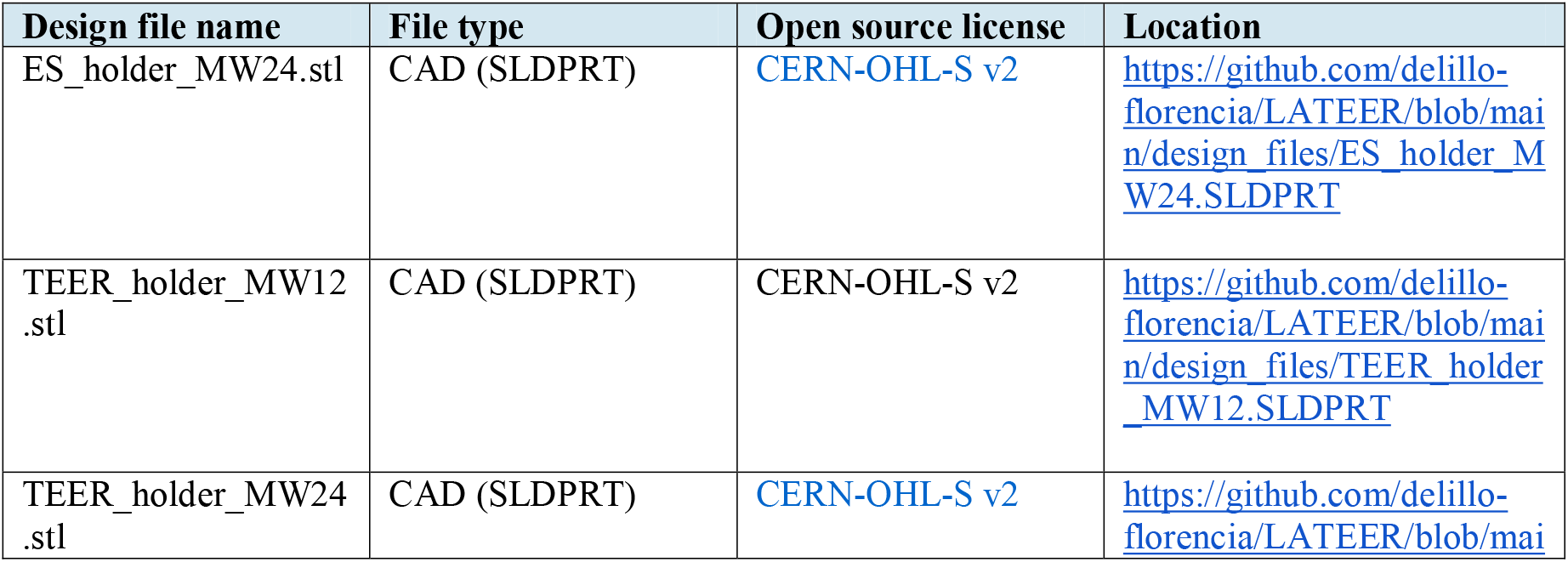

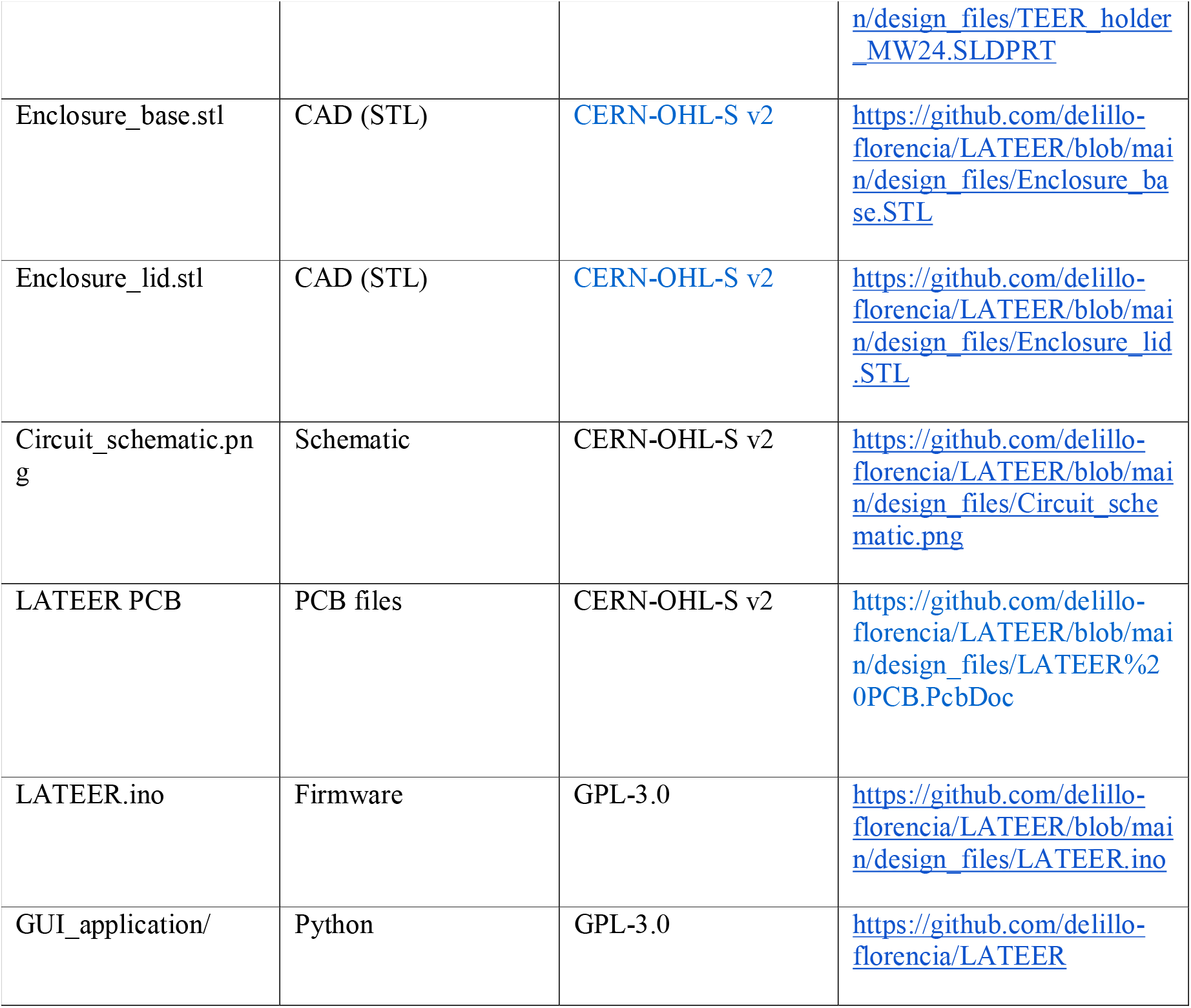

## Bill of materials summary

| Component | Qty | Cost/unit | Total | Source | Material |
| --- | --- | --- | --- | --- | --- |
| Arduino UNO | 1 | \$23 | \$23 | Arduino.cc / Local | Electronic |
| Custom PCB | 1 | \$27 | \$27 | Local PCB service | Electronic |
| Electronic components* | 1 set | \$22 | \$22 | DigiKey / Mouser | Electronic |
| 12V power supply | 1 | \$8 | \$8 | Local supplier | Electronic |
| PLA filament (100g) | 100g | \$2 | \$2 | 3D printing supplier | Polymer |
| Graphite leads 0.5mm 2H | 24 | \$0.08 | \$2 | Office supply store | Carbon |
| Metal terminals | 16 | \$0.50 | \$8 | Electronics store | Metal |
| Silicone adhesive | 1 | \$2 | \$2 | Hardware store | Polymer |
| Cables and connectors | 1 set | \$3 | \$3 | Electronics store | Electronic |
| **TOTAL** | | | **~\$97** | | |

\*Electronic components include: resistors, capacitors, LM317 voltage regulator, TL082 operational amplifier, transistors (2N2222), sockets, and misc. Detailed BOM available in the repository.

## Build instructions

### 5.1 3D printing

1. Download STL files from the repository
2. Print using PLA filament with standard settings (0.2 mm layer height, 20% infill)
3. Print time: approximately 1 hour to complete the holder set
4. Print the electronics enclosure base and lid separately

### 5.2 Electronics assembly

5 Manufacture PCB using the provided Gerber files (or order from PCB service)
6 Solder components following the schematic and component placement guide
7 Connect Arduino UNO to the PCB via header pins
8 Upload firmware using Arduino IDE
9 Verify circuit operation using an oscilloscope before biological use

### 5.3 Electrode holder assembly

10 Drill 0.5 mm holes in the culture plate lid following the holder template
11 Attach the 3D-printed holder to the plate lid using silicone adhesive
12 Insert larger metal terminals into the holder holes and heat-fix to PLA
13 Solder connection cables to terminals
14 Insert graphite leads through the smaller terminals
15 Connect the cable assembly to the main unit via the connector

### 5.4 Sterilization

16 Clean the electrode assembly with 70% ethanol
17 Place under UV light for 20-30 minutes
18 Allow to dry in a sterile environment before use

## Operation instructions

### 6.1 Software installation

1. Install Python 3.x with required libraries (PyQt5, pyserial, sqlite3)
2. Launch the GUI application from the repository
3. Connect the Arduino via USB and select the COM port in the interface

### 6.2 Electrical stimulation mode

4 Select ES mode and number of channels (1-4)
5 Select plate type (MW12 or MW24) -this affects field calculation
6 Configure parameters: amplitude (mV/mm), frequency (Hz), pulse width (ms), duration
7 Start stimulation; pause/resume as needed for media changes

### 6.3 TEER measurement mode

8 Select TEER mode and configure sampling interval and duration
9 Include a blank transwell (no cells) for baseline subtraction
10 Start measurement; data displayed in real-time
11 Export data to CSV for analysis

## Validation and characterization

### 7.1 Biocompatibility validation

Electrode biocompatibility was assessed using mesenchymal stem cells (MSCs), a cell type for which both electrical stimulation and TEER protocols are well established ^2,9,22^. Stainless steel electrodes coated with carbon ink, following the approach described by Jones and Chen, resulted in cell detachment and death within 48 hours of culture, accompanied by visible changes in medium color indicating pH shifts. In contrast, graphite electrodes showed no cytotoxic effects. MSCs maintained normal morphology and proliferated over 10 days of continuous electrode contact, with no observable changes in medium color **(Figure 2)**.

**Figure 2.**
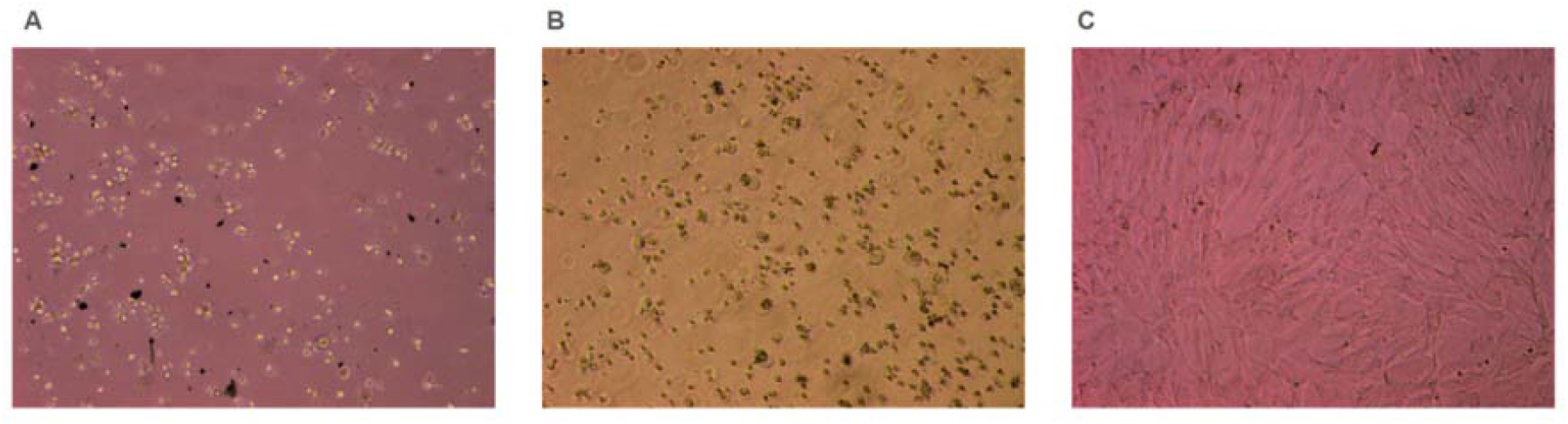
Results of the electrode biocompatibility test using MSCs after contact with the proposed materials. **(A)** Carbon rod electrodes. Although no change in the culture medium color was observed, cell death and material detachment were evident. **(B)** Stainless steel electrodes coated with five layers of graphite ink. A clear change in culture medium color was observed, indicative of pH alteration, accompanied by cell death.**(C)** Graphite pencil lead electrodes. Cell growth was not affected by the presence of graphite, with no observable changes in medium color and no evidence of cell death.

### 7.2 TEER measurements

To assess whether LATEER can resolve differences in cell coverage on a permeable support, human MSCs were seeded on 24-well transwell inserts at 20,000, 50,000 and 140,000 cells per insert, with a fourth, cell-free insert used as a blank. Nuclear staining with Hoechst confirmed attachment and the expected rank order of nuclear density across the three seeded inserts **(Figure 3 A–C)**. Graphite-lead electrodes were then mounted in the printed lid and the four inserts were recorded simultaneously with the four TEER channels of the device (1 reading every 5 s, ∼45 min).

**Figure 3.**
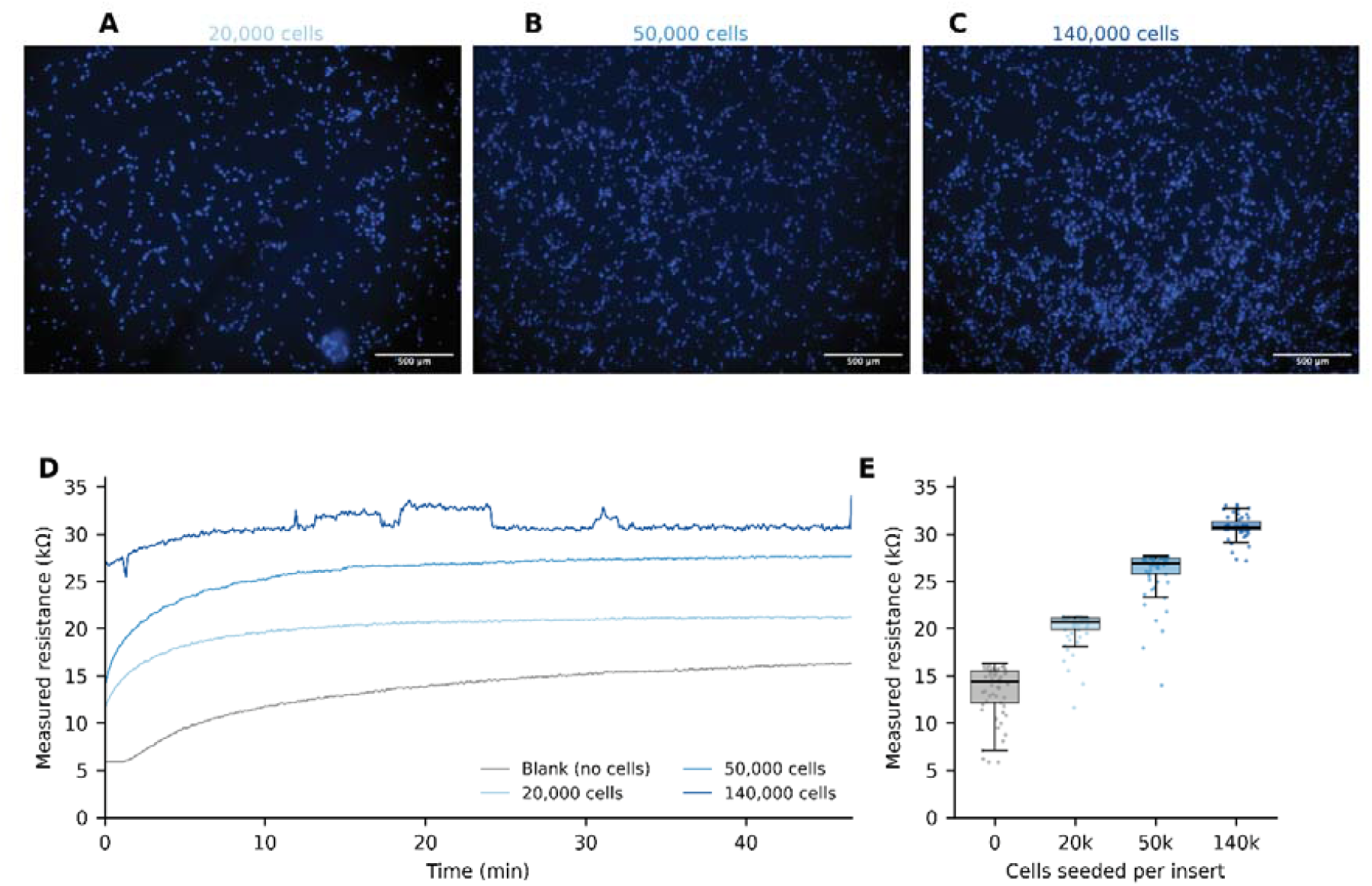
LATEER resolves cell-density differences in electrical resistance. (A–C) Hoechst-stained nuclei of human MSCs seeded on 24-well transwell inserts at (A) 20,000, (B) 50,000 and (C) 140,000 cells per insert, confirming attachment and the expected rank order of nuclear density; a fourth, cell-free insert served as a blank. (D) Resistance recorded simultaneously on the four TEER channels (one reading every 5 s, ≈45 min); all channels show a rising transient over the first ≈10 min and then stabilize. Traces follow a grey-to-dark-blue scale from the cell-free blank to the highest seeding density. (E) Box plots of the stable-phase readings with the individual measurements overlaid: median resistance increased monotonically with seeding number (blank 14.4 kΩ; 20,000 cells 20.8 kΩ; 50,000 cells 26.9 kΩ; 140,000 cells 30.7 kΩ). Because a single insert was recorded per condition, the overlaid points are repeated readings of the same well and not biological replicates, and no inferential statistic was applied. Values are raw two-electrode resistance (kΩ), not normalised TEER (Ω·cm^2^).

All four channels showed a rising transient during the first ∼10 min of recording, after which the readings stabilised (**Figure 3D)**. Once stable, the measured resistance increased monotonically with the number of cells seeded: 14.4 kΩ (median; IQR 12.2–15.5) for the cell-free insert, 20.8 kΩ (19.9– 21.1) at 20,000 cells, 26.9 kΩ (25.8–27.4) at 50,000 cells and 30.7 kΩ (30.4–31.3) at 140,000 cells **(Figure 3E)**. The resistance attributable to the cell layer (channel reading minus the blank insert) was therefore ∼6.6, ∼12.6 and ∼17.4 kΩ, respectively; the increase was monotonic but sublinear with respect to seeding number, consistent with a progressively more confluent layer.

Two limitations should be stated explicitly. First, a single insert was recorded per condition, so the points in Fig. 3 E are repeated readings of the same well rather than biological replicates, and no inferential statistics were applied. Second, the absolute resistance reported here is the raw two-electrode reading and is dominated by the electrode–medium path geometry rather than by the paracellular barrier itself; MSCs do not form a tight barrier, and the values are not converted to normalised TEER (Ω·cm^2^). Comparisons are therefore only meaningful between channels recorded simultaneously with a fixed electrode configuration. Consistently, the rank order was reproduced when the electrode lid was left in place across consecutive days, but absolute values shifted whenever the electrodes were remounted, which precludes quantitative comparison across days. Accordingly, this experiment supports the claim that the device discriminates cell-density-dependent differences in electrical resistance in real time, and not that it provides calibrated TEER values for a barrier-forming epithelium — the latter would require endothelial or epithelial monolayers with established TEER references.

The electronic accuracy underlying these readings was validated independently against commercial resistors and a potentiometer connected to the four channels in parallel **(Figure S3)**: measured values agreed with nominal values to within 5% for R □ 4.7 kΩ, with larger relative error at lower resistances due to a systematic additive offset (∼+200 Ω); the four channels tracked a continuously varied resistance with small inter-channel differences **(Figure S3)**, confirming that the resistance changes reported above are within the validated operating range of the instrument.

### 7.3 Electrical stimulation of cardiomyocytes

To functionally validate the electrical stimulation capabilities of the developed platform, two complementary proof-of-concept experiments were performed on stem cell-derived cardiomyocytes differentiated following the Lian et al. protocol ^23^: (i) an exploratory experiment that identified the field-strength capture threshold and screened frequency and pulse-width effects (figures below), and (ii) a systematic parameter sweep on lactate-selected human embryonic stem cell-derived cardiomyocytes confirming reproducible capture.

While MSCs were suitable for biocompatibility assessment due to their widespread use in electrical stimulation studies and well-characterized responses to electrode contact, hESC-CMs provide distinct advantages for validating stimulation functionality. Cardiomyocytes exhibit spontaneous rhythmic contractions that can be captured and paced by external electrical fields, enabling direct visual confirmation of successful stimulation through changes in beating frequency. This frequency capture phenomenon—where cells synchronize their intrinsic rhythm to the applied stimulation frequency— serves as an unambiguous functional readout that is not achievable with non-excitable cell types.

#### Experimental protocol and frequency capture assessment

Cardiomyocyte monolayers were subjected to external electrical pacing using the proposed stimulation system while contractile activity was recorded by video microscopy. Each recording followed a standardized 60-second protocol consisting of three consecutive 20-second phases: baseline spontaneous contraction, active electrical stimulation, and post-stimulation recovery. This 20-20-20 protocol enables direct comparison of contractile parameters across phases within the same recording, minimizing inter-sample variability. Contractile velocity was quantitatively extracted from video recordings using ContractionWave software.

To test out the device, we first sought to modulate the signal amplitude between sub-threshold and successful frequency capture conditions. At 250 mV/mm (**Figure 4A, Video S1**), the contraction pattern during the stimulation phase (green) remains indistinguishable from the baseline phase (black), indicating that this field strength is insufficient to induce membrane depolarization. In contrast, at 400 mV/mm (**Figure 4B, Video S2**), contractions during the stimulation phase become visibly regularized and synchronized to the applied frequency, demonstrating successful frequency capture. The recovery phase (blue) shows the return of spontaneous activity following stimulation cessation.

**Figure 4.**
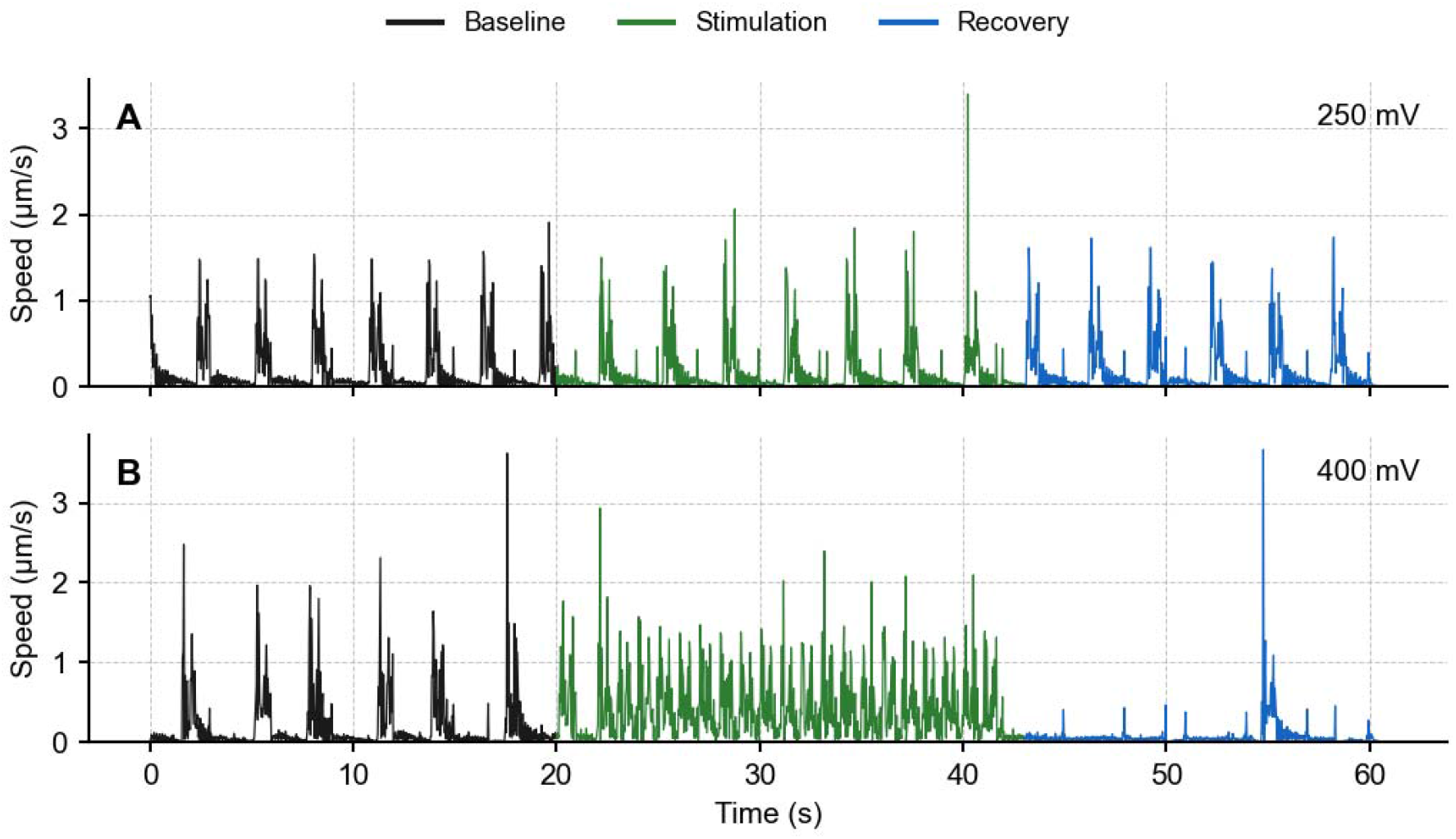
Amplitude-dependent frequency capture of stem cell-derived cardiomyocytes. Contraction velocity was extracted from video microscopy (ContractionWave) over the standardized 60-s, three-phase 20–20–20 s protocol: baseline spontaneous activity (black), electrical stimulation (green) and post-stimulation recovery (blue). (A) At 250 mV/mm (Video S1) the stimulation-phase trace is indistinguishable from baseline, indicating a sub-threshold field. (B) At 400 mV/mm (Video S2) contractions during stimulation become regularized and synchronized to the applied pacing signal, demonstrating successful capture; spontaneous activity resumes during recovery. These recordings correspond to the exploratory stimulation experiment described in Section 7.3.

#### Frequency-domain analysis and parameter sweeps

To quantitatively assess frequency capture across additional parameter combinations, Fast Fourier Transform (FFT) analysis was performed on the contraction velocity signals. FFT provides an objective metric: when successful pacing occurs, the dominant spectral peak shifts from the intrinsic beating rate to a frequency corresponding to the applied stimulation.

FFT spectra computed on the 20-second stimulation window (**Figure 5A**) demonstrate that once capture is achieved at fixed field strength and pulse width, stimulation at 1 Hz and 2 Hz produced dominant velocity-peak frequencies of 2.00 and 3.99 Hz respectively (±0.05 Hz; 400 mV/mm, 6 ms), confirming synchronization to the applied frequency (**Panel B**). Pulse width had minimal effect on capture efficiency above threshold; sweeping from 2 ms to 100 ms at fixed amplitude (400 mV/mm) and frequency (1 Hz) consistently yielded the same dominant frequency (**Panel C**). Because capture is an all-or-none response, this establishes only that all widths tested were supra-threshold and does not imply a graded effect on capture efficacy. It does carry a practical consequence: at 1 Hz the charge injected per cycle differs roughly fiftyfold between a 2 ms and a 100 ms pulse for no measurable benefit, so the shortest effective pulse should be preferred in order to limit faradaic product formation and electrode wear.

**Figure 5.**
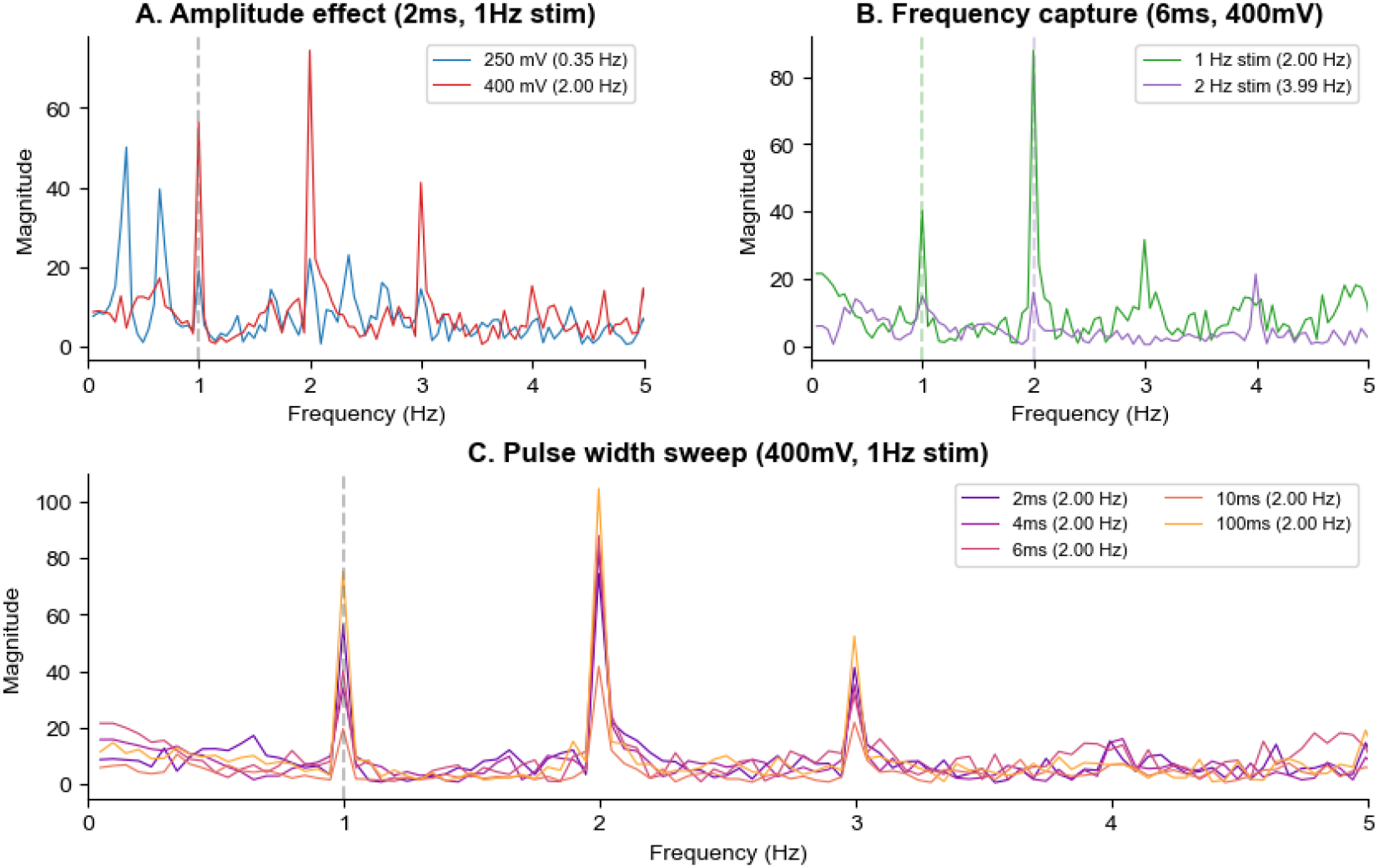
Frequency-domain confirmation of pacing across stimulation parameters. Fast Fourier Transform (FFT) of the contraction-velocity signal computed on the 20-s stimulation window. (A) Representative FFT spectra under successful capture, showing a dominant peak locked to the applied pacing; the peak appears at twice the stimulation frequency because each pulse produces one contraction–relaxation cycle, i.e. two velocity peaks per period, with additional harmonics arising from the impulse-like contraction waveform. (B) Varying stimulation frequency at fixed amplitude (400 mV/mm) and pulse width (6 ms): 1 Hz and 2 Hz pacing yield dominant peaks at 2.00 Hz and 3.99 Hz, respectively. (C) Varying pulse width from 2 to 100 ms at fixed amplitude (400 mV/mm) and frequency (1 Hz) leaves the dominant frequency unchanged, indicating that pulse width has minimal effect on capture once threshold is exceeded. The corresponding time–frequency evolution across all three phases is shown in the spectrogram of Figure S4.

The dominant peaks in successful capture conditions appear at twice the stimulation frequency (e.g., 200 Hz when stimulating at 1 Hz), along with additional harmonic content. This doubling arises from the mechanical nature of the contractile signal: each stimulation pulse triggers a contraction-relaxation cycle producing two velocity peaks per period. The non-sinusoidal, impulse-like character of cardiomyocyte contractions further distributes energy across harmonics when decomposed through Fourier analysis. The doubled peak is therefore an artefact of signal rectification and must not be read as 2:1 capture or as alternans; 1:1 correspondence between pulses and contractions is confirmed directly in the time-domain traces and videos

The full 60-s recording is also summarized as a time–frequency spectrogram: **Figure S4** shows the dominant contraction-frequency band held at the spontaneous rate (≈0.25–0.35 Hz) during baseline, shifting to the applied frequency at stimulation onset (t = 20 s) and returning to spontaneous activity after cessation, confirming that the spectral shifts during stimulation are attributable to external pacing rather than to analysis artifacts.

The device was also tested on lactate-selected cardiomyocytes, where reliable frequency capture was likewise achieved. Notably, in several fields the device paced regions of the monolayer that were quiescent at baseline, driving contraction where none was present before stimulation (Video S3), further confirming that the observed activity was externally imposed rather than spontaneous.

#### Capabilities and limitations of LATEER

- Dual electrical-stimulation and TEER operation from a single open-source unit built for < USD 100, with four independent channels and a plain-language GUI.
- Configurable pulsatile stimulation (amplitude up to 8.2 V, 0.1–500 Hz, pulse width ≥ 0.1 ms) that reliably paces stem cell-derived cardiomyocytes; field-strength capture threshold in the 250–400 mV/mm range.
- Resistance measurement from 300 Ω to 1 MΩ (<5% error for R ⊡ 4.7 kΩ) across four channels; discriminates cell-density-dependent resistance differences in real time.
- Zero-fabrication graphite pencil-lead electrodes (≈ USD 2) that are biocompatible over ≥ 10 days and, with open-ring 3D-printed holders, allow microscopy without removing the electrodes.
- Limitations: monophasic (DC) rather than AC drive; TEER values are raw two-electrode resistance (kΩ), not normalised TEER (Ω·cm^2^), and are comparable only between channels recorded simultaneously; absolute values shift on electrode remounting, precluding cross-day comparison. Biological validation used n = 1 insert per condition (no biological replicates) and non-barrier-forming MSCs, so calibrated barrier TEER remains to be demonstrated with endothelial/epithelial monolayers.

## Conclusions

LATEER demonstrates that electrical stimulation and TEER sensing, normally provided by two separate commercial instruments costing USD 2,500–9,000, can be delivered by a single open-source device for under USD 100. The device paces stem cell-derived cardiomyocyte monolayers reproducibly, capturing contraction at field strengths above the 250–400 mV/mm across the 0.5–5 Hz range, and resolves cell-density-dependent differences in electrical resistance in real time, discriminating a sevenfold range of seeding densities on the same plate. Because the electrodes remain fixed in the plate lid and only a single cable enters the incubator, whichever function is in use operates continuously under normal culture conditions, removing the temperature excursions and placement variability that dominate the error budget of manual measurement, while the open-ring holders keep the monolayer visible to the microscope throughout.

The capabilities demonstrated here define a considerably larger application space than the one tested. The validated stimulation envelope — 0.1–500 Hz, pulse widths from 0.1 ms and field strengths up to 600 mV/mm — encompasses the parameter ranges used in published protocols for electrical conditioning of hPSC-CMs, including the low-frequency, millisecond-pulse regimes associated with improved sarcomere alignment, calcium handling and conduction velocity, and the progressive frequency-ramping schemes used in intensity training ^8,24^. Combined with four independently addressable channels and uninterrupted in-incubator operation, the device is in principle capable of running the multi-week conditioning schedules such protocols require, with several parameter arms in parallel on a single plate. Whether it drives maturation has not been tested here and cannot be inferred from pacing alone.

Stimulation and resistance measurement presently require different electrode geometries and are therefore run as separate experiments. Consolidating them into a single holder, with the two modes interleaved in firmware between stimulation pulses, is a tractable extension of the existing design and would enable a measurement that neither class of commercial instrument currently offers: label-free resistance monitoring of the same monolayer across a conditioning protocol, so that the electrical signature of a culture is followed as a longitudinal readout rather than sampled at endpoints.

By pairing zero-fabrication pencil-lead electrodes with microscopy-compatible holders and freely available design files, firmware and software (CERN-OHL-S v2), LATEER lowers the cost and technical barrier to combined electrophysiological readout and stimulation, particularly for resource-limited laboratories. Future work will address alternating-current operation and calibrated barrier-TEER validation with endothelial or epithelial monolayers.

## Ethics statements

This work did not involve human subjects or animal experiments. All cell lines used were established, previously characterized lines. H1 hESC was obtained from WiCell. Human mesenchymal stromal cells were obtained from human umbilical cord Wharton’s Jelly. Human umbilical cord was obtained via written informed consent obtained from each mother before normal cesarean birth. The donation is anonymous and the human umbilical cords were obtained from discarded placentas. All experiments and methods were performed in accordance with relevant guidelines and regulations. All experimental protocols and informed consents were approved by the FLENI Ethics Committee.

## CRediT authorship contribution statement

**Florencia De Lillo:** Conceptualization, Methodology, Software, Hardware, Investigation, Validation, Writing – original draft. **Joaquin Smucler:** Conceptualization, Methodology, Hardware, Resources (cell culture), Supervision, Writing – review & editing.

## Acknowledgments

The authors thank the LIAN laboratory at Fundación FLENI for providing facilities and support for cell culture experiments.

