## SUPPLEMENTAL MATERIAL for "LATEER: Low-Cost Open-Source Platform for Electrical Stimulation and TEER Measurement in Human Cardiomyocytes"

**Supplemental Tables**

***Table S1:*** *COMSOL Multiphysics® simulation parameters, materials, and assumptions*

| **Category** | **Parameter** | **Description / Value** |
| --- | --- | --- |
| **Physics regime** | Study type | Stationary (electrostatic) |
|  | Governing equation | Electric field defined as **E = −∇V** |
|  | Coordinate system | Global Cartesian coordinate system |
| **Geometry** | Geometry design | Electrode and culture well geometries modeled in SolidWorks and imported into COMSOL |
|  | Well diameter | 2.5 cm |
|  | Monolayer height | 1 mm |
| **Electrode materials** | Graphite electrodes | COMSOL built-in *Graphite sheet [solid]* material |
|  | Platinum electrodes (L-shaped reference) | COMSOL built-in Platinum material |
|  | Carbon rod electrodes | COMSOL built-in *Carbon* material |
| **Culture medium** | Medium type | Dulbecco’s Modified Eagle Medium (DMEM) |
|  | Electrical conductivity | 1.54 S/m |
| **Cell monolayer model** | Representation | Cylindrical domain corresponding to the culture area |
|  | Effective resistivity | ~302 Ω·m |
|  | Source of resistivity value | Derived from reported TEER measurements of endothelial cell monolayers in the literature |
| **Electrical stimulation parameters** | Applied voltage | 5V |
| **Analysis output** | Evaluated quantity | Electric field magnitude and spatial distribution |
| **Model assumptions** | Material behavior | All materials assumed linear, homogeneous, and isotropic |
|  | Electrochemical effects | Not included (purely electrostatic approximation) |
|  | Field–distance relationship | Inverse relationship between electric field magnitude and electrode separation |

**Supplemental Videos**

**Video S1:** 250 mV/mm stimulation of non-purified hESC-CMs without capture

**Video S2:** 400 mV/mm stimulation of non-purified hESC-CMs with capture

**Video S3:** Contraction induction on purified, non-beating hESC-CMs

**Supplemental Figures**

**Figure S1:** LATEER electrode and well-plate configuration. A) Dual metal terminal system for secure graphite lead attachment. B) Lateral view of the electrode and well-plate arrangement. C) TEER measuring configuration. D) Electrical stimulation configuration.
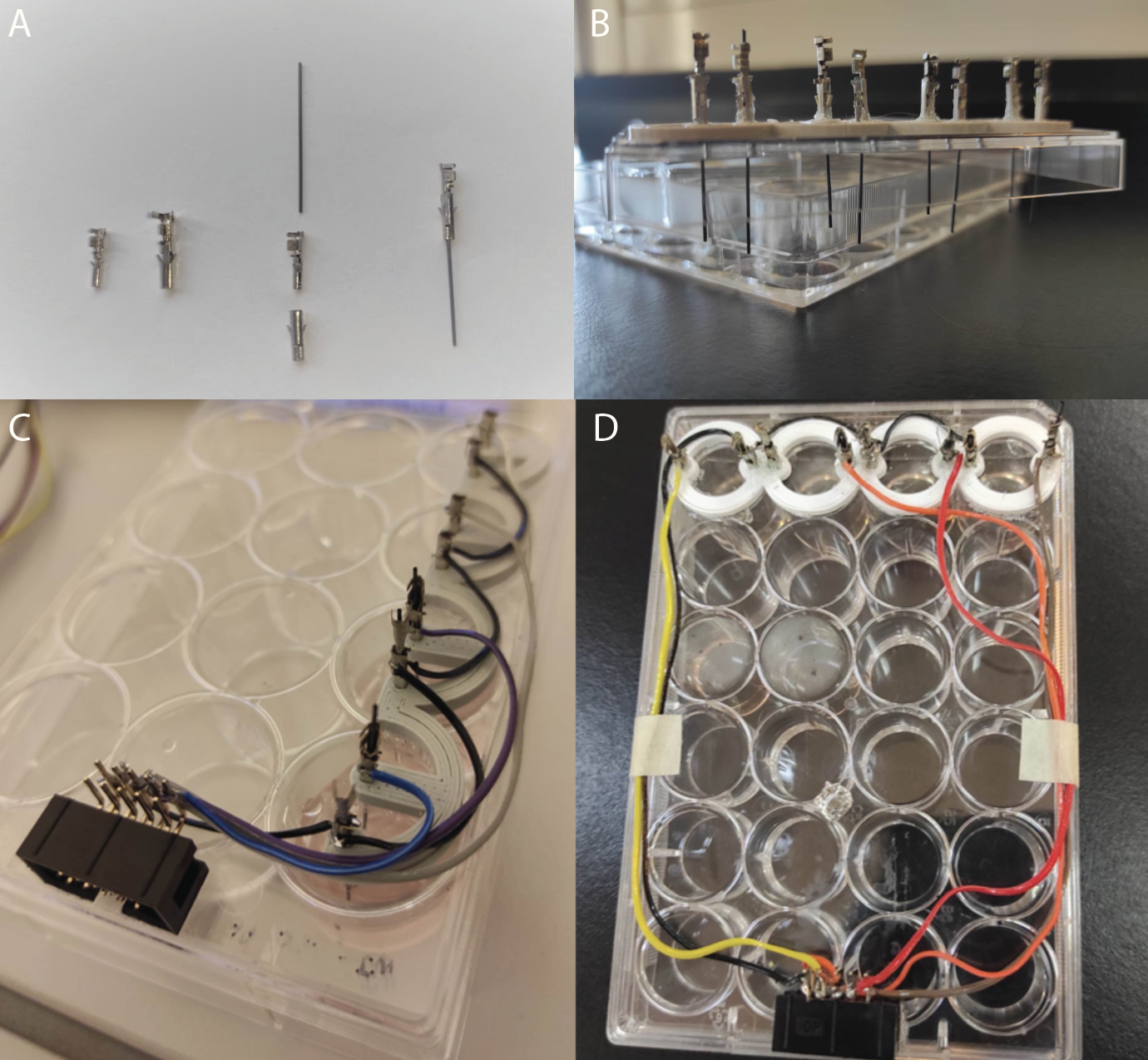

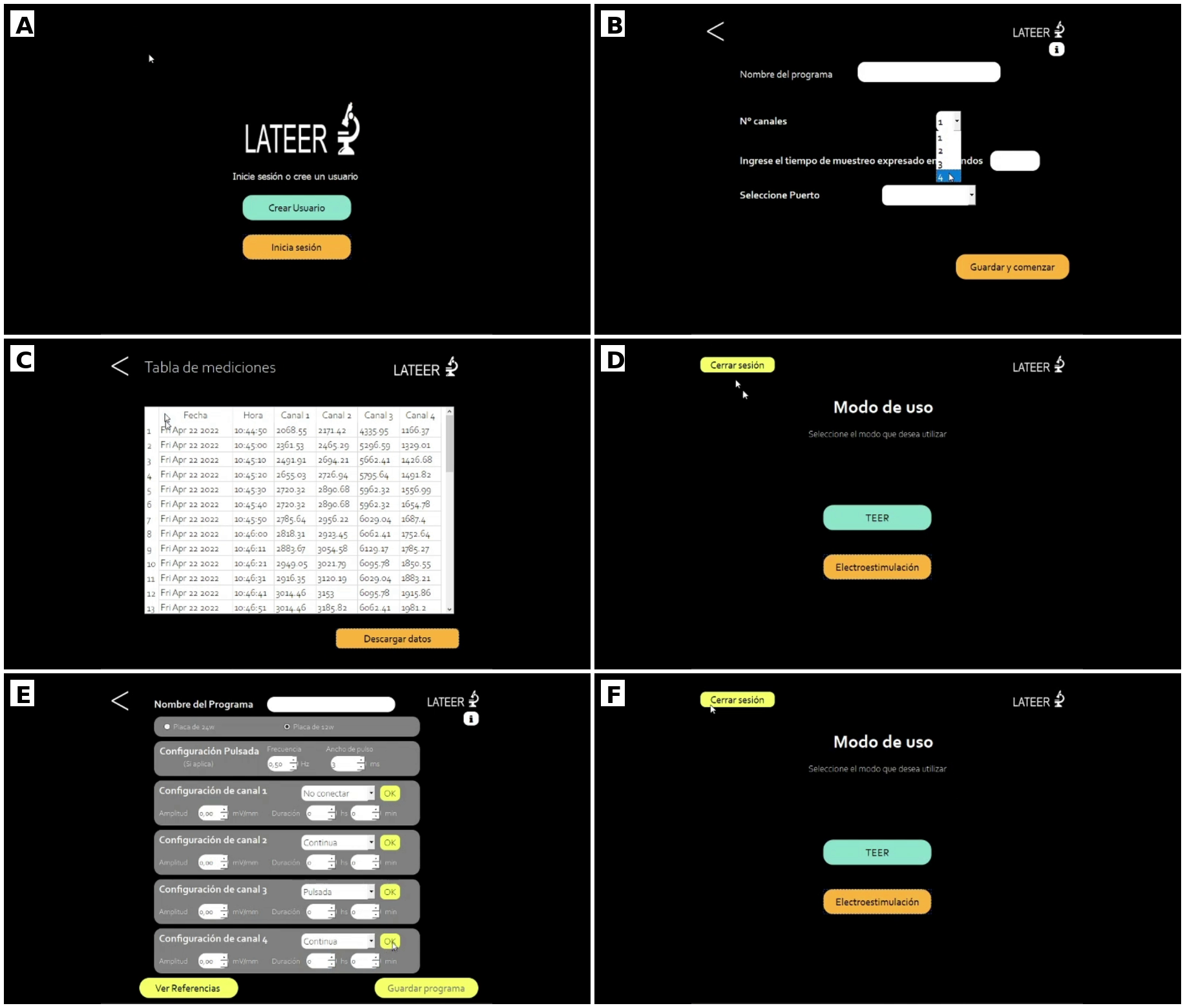

**Figure S2:** LATEER graphical user interface. (A) Login / user-account screen. (B) New-program setup: program name, number of channels, sampling interval, and serial-port selection. (C) TEER results table with per-channel resistance readings and CSV export. (D) Mode-selection screen (TEER or Electroestimulación) reached after login. (E) Electrical-stimulation program configuration: plate format (12- or 24-well), pulsed waveform frequency/pulse width, and independent per-channel amplitude, connection mode, and duration. (F) Saved-programs / mode-selection navigation. All screens use plain-language labels and require no coding or electronics knowledge to operate.

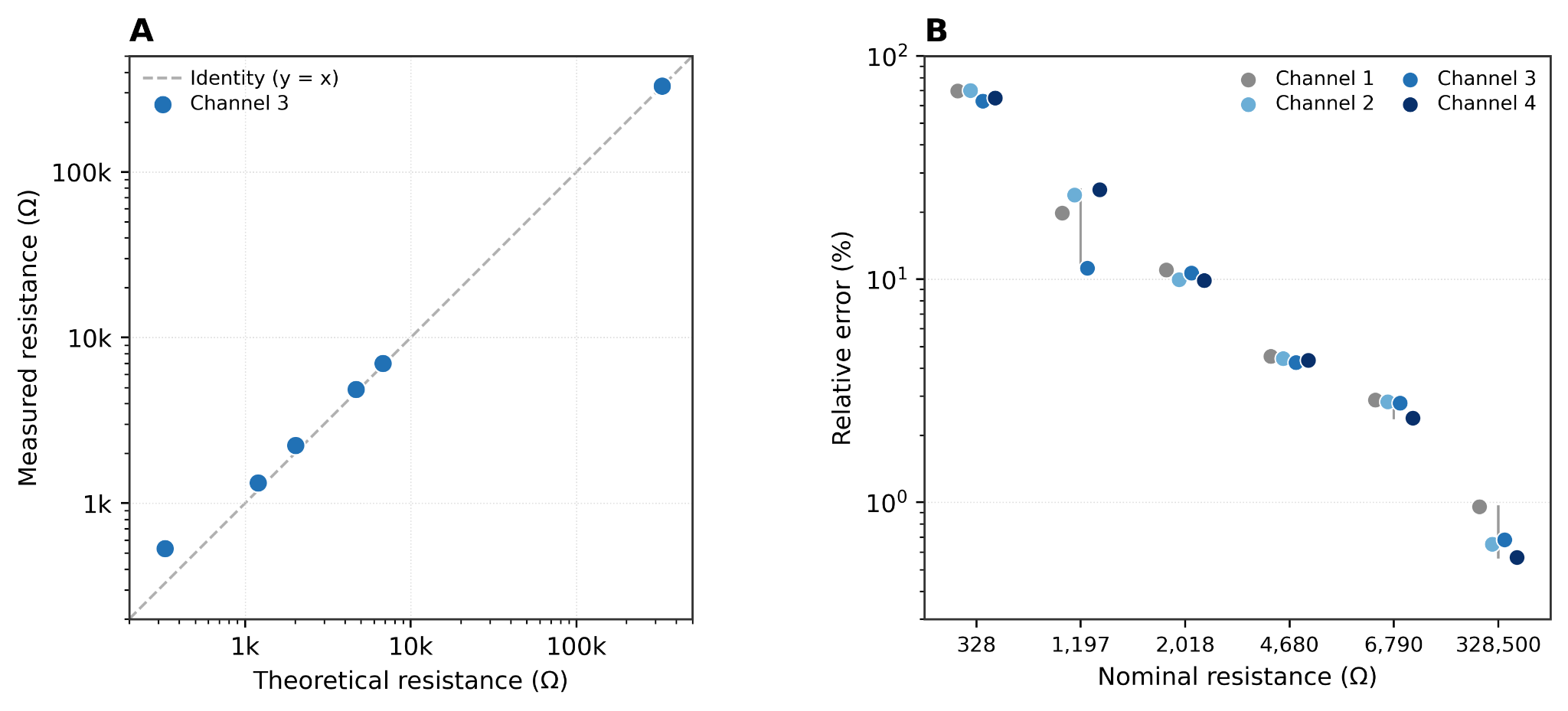

**Figure S3:** Electronic validation of the four TEER channels against commercial resistors. (A) Measured versus theoretical resistance for Channel 3 (lowest mean relative error across the validated range), spanning 328 Ω to 328.5 kΩ; dashed line, identity (y = x). (B) Relative error (%, |measured − nominal| / nominal) for all four channels at each nominal resistance value; grey bars indicate the min–max spread across channels. Error falls below 5% for R ≳ 4.7 kΩ and is dominated by a systematic additive offset (~+200 Ω) at lower resistances. Channels show consistent performance except at the 1,197 Ω point, where Channel 3 was measured against a nominally different resistor than Channels 1, 2 and 4 (985–998 Ω).

**Figure S4:** The spectrogram representation visualizes the temporal evolution of frequency content across the three experimental phases. A clear transition in the dominant frequency band occurs at stimulation onset (t = 20 s), shifting from spontaneous rhythm to the applied frequency. At stimulation cessation (t = 40 s), a transient period of reduced contractile activity precedes the gradual recovery of spontaneous contractions. This post-stimulation quiescent interval is consistent with a transient refractory phase commonly observed following paced excitation of cardiomyocyte monolayers.
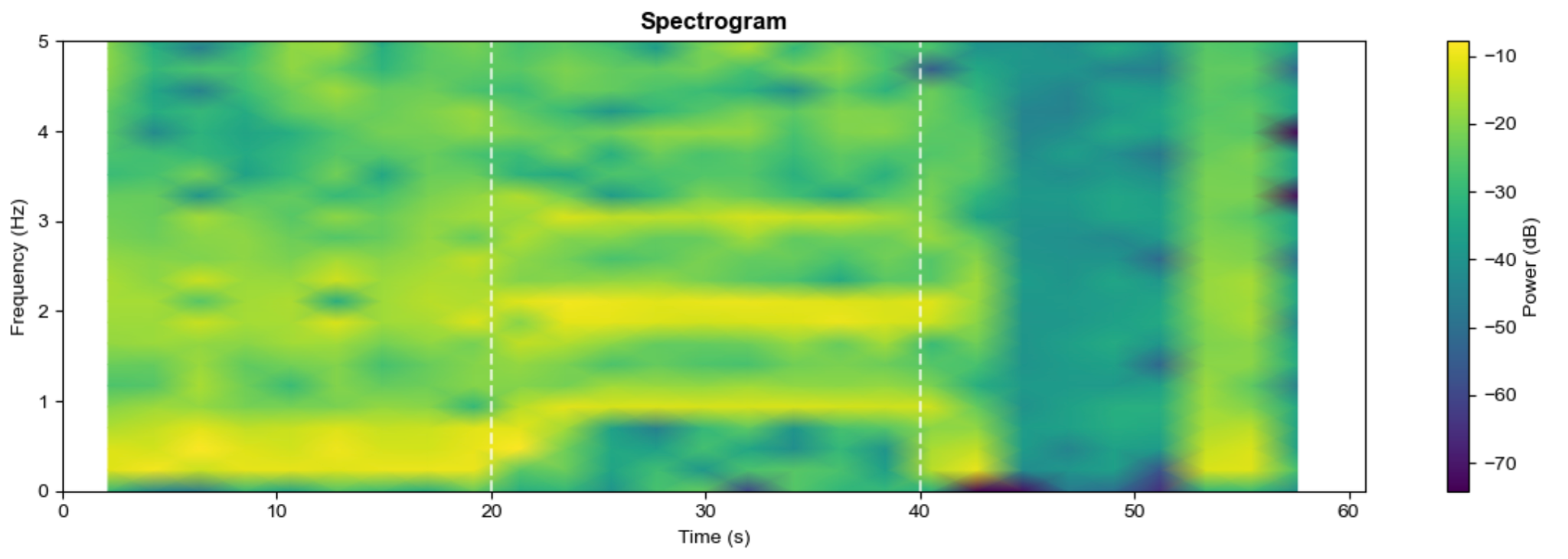
